# MacroMove: a global database of animal movement across taxa and movement types

**DOI:** 10.64898/2025.12.08.692918

**Authors:** Caitlin Wilkinson, Emilio Berti, Alexander Dyer, Myles H. M. Menz, Carsten Meyer, Marlee A. Tucker, Kate L. Wootton, Maarten J.E. Broekman, Selwyn Hoeks, Jana Roth, Remo Ryser, Myriam R. Hirt

## Abstract

Animal movement is a key ecological process that shapes species distributions and interactions and connects resources, populations and habitats, underpinning ecosystem functioning and stability. While foraging, dispersal, and migration are the most widely studied movement types, they operate at distinct spatial and temporal scales, and no single resource currently integrates these processes across taxa, traits, and ecosystems. We present MacroMove, a comprehensive global movement database containing 92,207 observations from 4,329 (4,167 using GBIF backbone) species, spanning 36 classes and 648 families across 205 countries, with data sourced from 6,234 references and 36 meta-analyses/databases. It includes both vertebrates and invertebrates from terrestrial, freshwater, and marine environments. The database includes various movement metrics - home range area, dispersal distance, migration distance, day range, routine speed and maximum speed - including summary statistics (e.g. mean, median, maximum) where available. Records are reported at individual, population, or species levels and include standardised metadata describing species identity (binomial name, class, family), research methods (e.g. sampling method, sampling level and sampling dates) and location (e.g. country, location and coordinates). The dataset was assembled from published meta-analyses/databases and original studies included in these meta-analyses. Trait information (body mass and locomotion mode) from original studies and associated metadata was incorporated, with additional data sourced from established trait databases and reputable online resources. In the final dataset, body mass is available for 98 % and locomotion for 100% of species. Taxonomy was harmonised using the Global Biodiversity Information Facility (GBIF) backbone taxonomy. The database is supplied in comma-delimited text (.csv) format to maximise accessibility and compatibility with a wide range of workflows. By consolidating existing movement data into a single, standardised and openly accessible resource, MacroMove fills a major gap in available biodiversity data. It serves as a practical and extensible foundation for synthesising movement information across taxa and ecosystems, advancing data-driven research on the ecological and evolutionary correlates of animal movement.

## Open Research Statement

### Introduction

Movement is a ubiquitous process shaping biodiversity patterns across spatiotemporal scales (Jeltsch et al. 2013). The way species move drives species interactions (McIntyre & Wiens. 1999; Schweiger et al. 2012), community composition (MacArthur & Wilson. 1967; Leibold et al. 2004), the spatial distribution of species (Pulliam. 2000) and changes in species’ traits and genetic diversity (e.g. Bruggeman et al. 2010; Cushman & Lewis. 2010; Shafer et al. 2012;). Anthropogenic habitat fragmentation, land-use changes, biological invasions and climate change alter ecosystems affecting how species respond to these disturbances (Nathan et al. 2008). As the current global biodiversity crisis has major consequences for ecosystem stability, functioning and services (Cardinale et al. 2012; Harrison et al. 2014; Pennekamp et al. 2018), it is critical to understand how species will move and adapt to global changes.

The three most common types of movement are foraging, dispersal and migration, which occur at distinct spatiotemporal scales and can couple different habitat types (Clobert et al. 2012; Jeltsch et al. 2013; Guzman et al. 2019). Foraging (or exploratory) movement describes how species search for and exploit resources within their home range. It can occur several times per day and structures local habitats, species interactions and community composition (Pawar et al. 2012; Jeltsch et al. 2013; Hirt et al. 2017; Guzman et al. 2019). Foraging can occur over varying spatial scales depending on species body size and trophic level, where smaller species remain localised and larger species move across multiple habitats in a region to meet their resource requirements (McCann et al. 2005; McCauley et al. 2012; Guzman et al. 2019). Dispersal is the long-term permanent movement of individuals to a new site of reproduction, occurring at any life stage and at any spatial scale above an individual’s home range (Bowler & Benton. 2005; Clobert et al. 2009). It is a unique event in a species lifetime and can take a considerable amount of time when compared to the other movement types (Jeltsch et al. 2013). Additionally, dispersal takes place within heterogeneous landscapes as a response to population density or suboptimal conditions and results in colonisation of novel habitats, increased species ranges and connection of populations across habitats (Holt & Keitt. 2005; Baguette et al. 2013; Matthysen. 2012; Guzman et al. 2019). Migration is the cyclical tracking of available resources or mates resulting in a return to the origin site, arising once or at regular seasonal intervals and completed over varying durations (e.g., days, months or years) (Dingle. 1996; Jeltsch et al. 2013). Migratory movements tend to be of greater scale and length than the other movement types and have higher predictability due to proximate cues of body condition, climate and phenology (Dingle & Drake. 2007; Guzman et al. 2019). Due to the round-trip nature of migration, it does not connect different populations but allows the movement of single populations through many different habitats (Guzman et al. 2019). Movement speed affects all movement behaviours, determining their intensity, duration and likelihood of success (Wilson et al. 2015). For example, maximum speeds shape predator–prey dynamics by influencing the outcome of feeding or hunting movements (Wilson et al. 2013), whereas routine travel speeds can reflect an animal’s realised movement capacities across landscapes (Dyer et al. 2023).

A range of methods and metrics (e.g. area, distance and speed) are used to quantify animal movement. Area is a way to measure animal space use, which is most commonly assessed as home range size (Burt 1943). There are several extensive datasets that report home range size for marine and terrestrial mammals (e.g. Tucker et al. 2014; Broekman et al. 2023) and for other vertebrates (e.g. McCauley et al. 2015; Tamburello et al. 2015). Distance traveled is a key ecological metric that quantifies connections between animal behavior, energy use, and demographic processes, providing valuable insights into species’ movement patterns and ecological dynamics (Rowcliffe et al. 2012). Technological advances of GPS and other methods have allowed detailed tracking of species movement and distances between locations (Morales et al. 2010). Various meta-analyses have collated distance data for different movement processes, such as daily movement distance (or day range) in mammals (Carbone et al. 2005), dispersal distance for birds (e.g. Bowman et al. 2011), mammals (e.g. Whitmee & Orme. 2013), amphibians (e.g. Smith & Green. 2005), fish (e.g. Comte & Olden. 2018) and insects (e.g. Komonen & Muller. 2018) and also migration distance for reptiles (e.g. Hays & Scott, 2013), mammals (e.g. Joly et al. 2019), amphibians (e.g. Trochet et al. 2014) and both vertebrates and invertebrates (e.g. Hein et al. 2012). Over the last century, there have been major advancements on measuring speed, with compiled datasets on maximum speeds (Hirt et al. 2017) and routine (Dyer et al. 2023) and locomotor performance speeds (Cloyed & Dell. 2020).

Despite the extensive study of animal movement, we lack a comprehensive database that encompasses foraging, dispersal and migration, across metrics, taxa, traits and ecosystem types. We have compiled a unified movement database that consolidates available data on animal home range areas, dispersal and migration distances, day ranges, and movement speeds across mammals, fishes, birds, reptiles, invertebrates. Importantly, these movement records include metadata providing additional context, such as temporal and spatial information (e.g., country, study duration, start and end coordinates) and general species traits (e.g. body mass and locomotion mode).

## METADATA

### Class I. Data set descriptors

**A. Dataset identity:**
Title: MacroMove: a global database of animal movement across taxa and movement types
**B. Dataset and metadata identification codes:**
MacroMove_db.csv MacroMove_refs.csv MacroMove_taxonomy.csv MacroMove_Metadata
**C. Dataset description**

**1. Originators:**
Caitlin Wilkinson. German Centre for Integrative Biodiversity Research (iDiv) Halle-Jena-Leipzig, Theory in Biodiversity Science,Puschstraße 4, 04103, Leipzig; Friedrich-Schiller-Universitat Jena, Fürstengraben 1, 07743, Jena, Germany.
Myriam R. Hirt. German Centre for Integrative Biodiversity Research (iDiv) Halle-Jena-Leipzig, Theory in Biodiversity Science,Puschstraße 4, 04103, Leipzig; Friedrich-Schiller-Universitat Jena, Fürstengraben 1, 07743, Jena, Germany.
Remo Ryser. Institute for Plant Sciences (IPS), Bern, University of Bern, Altenbergrain 21, 3013, Bern, Switzerland.
**2. Abstract:**
Animal movement is a key ecological process that shapes species distributions and interactions and connects resources, populations and habitats, underpinning ecosystem functioning and stability. While foraging, dispersal, and migration are the most widely studied movement types, they operate at distinct spatial and temporal scales, and no single resource currently integrates these processes across taxa, traits, and ecosystems. We present MacroMove, a comprehensive global movement database containing 92,207 observations from 4,329 (4,167 using GBIF backbone) species, spanning 36 classes and 648 families across 205 countries, with data sourced from 6,234 references and 36 meta-analyses/databases. It includes both vertebrates and invertebrates from terrestrial, freshwater, and marine environments. The database includes various movement metrics - home range area, dispersal distance, migration distance, day range, routine speed and maximum speed - including summary statistics (e.g. mean, median, maximum) where available. Records are reported at individual, population, or species levels and include standardised metadata describing species identity (binomial name, class, family), research methods (e.g. sampling method, sampling level and sampling dates) and location (e.g. country, location and coordinates). The dataset was assembled from published meta-analyses/databases and original studies included in these meta-analyses. Trait information (body mass and locomotion mode) from original studies and associated metadata was incorporated, with additional data sourced from established trait databases and reputable online resources. In the final dataset, body mass is available for 98 % and locomotion for 100% of species. Taxonomy was harmonised using the Global Biodiversity Information Facility (GBIF) backbone taxonomy. The database is supplied in comma-delimited text (.csv) format to maximise accessibility and compatibility with a wide range of workflows. By consolidating existing movement data into a single, standardised and openly accessible resource, MacroMove fills a major gap in available biodiversity data. It serves as a practical and extensible foundation for synthesising movement information across taxa and ecosystems, advancing data-driven research on the ecological and evolutionary correlates of animal movement.
**D. Keywords/phrases:**
Animal movement, foraging, dispersal, migration, home range, movement speed, body mass, locomotion mode, vertebrates, invertebrates
**E. Description:**
The MacroMove database integrates movement observations with broad coverage across taxa, regions, and movement metrics (Table 1). The database currently contains 92,207 observations from 4,329 (4,167 using GBIF backbone) species, spanning 36 classes and 648 families across 205 countries, with data sourced from 6,234 references and 36 meta-analyses/databases. Geographic coordinates are available for 90% of movement data. After harmonisation and trait annotation, trait coverage consists of 98% for body mass and 100% for locomotion mode. Overall, 99.92% of the database (n = 92,133) comprises animal observations, while 0.08% (n = 74) represents microorganism data. Hereafter, all reported species numbers refer exclusively to those harmonised using the GBIF backbone.
The area component comprises home range records (Table 1). Home range is the sole area category, with 80,416 observations from 1,906 species across 16 classes, 363 families, and 158 countries, supported by 3,925 references. Trait completeness for area records is exceptionally high, with high trait coverage of 98% - 100%, and geographic coordinates for 97%.
The distance component comprises dispersal, migration, and day range records (Table 1). Dispersal is the largest distance category, with 6,868 observations from 1,853 species across 15 classes, 371 families, and 92 countries, supported by 1,418 references. Migration contributes 1,132 observations from 720 species, spanning 10 classes, 163 families, and 92 countries, based on 519 references. Day range includes 233 observations from 205 mammal species, representing 44 families across 20 countries, derived from 58 references. For distance records, trait completeness remains high (91–100%), whereas coordinate availability ranges from 6–56%.
The speed component includes routine and maximum speed records (Table 1). Routine speed comprises 1,603 observations from 718 species, across 26 classes and 238 families in 58 countries, drawn from 288 references. Maximum speed contributes 1,955 observations from 729 species, representing 21 classes, 216 families, and 32 countries, supported by 242 references. Trait coverage for speed records is likewise high (99–100%), while coordinate availability ranges 3–49%.
Together, this database forms a globally distributed resource spanning 212 countries and ∼8 orders of magnitude in body mass, encompassing broad taxonomic and ecological diversity across vertebrates and invertebrates. Figures 1–3 show the global distribution of movement observations, the number of countries contributing data, and the number of observations by taxonomic category, respectively.

**Table 1.** Overview of movement observations contained in the MacroMove database. For each movement metric, we provide the total number of entries, species (using GBIF backbone), taxonomic classes and families, countries represented, and original references (Refs). We also report data completeness for body mass, locomotion mode, and geographic coordinates. **Note:** The number of entries may exceed the number of independent measurements because some meta-analyses/databases report both species-level averages and individual observations.

| **Movement metric** | **Entries** | **Species** | **Classes** | **Families** | **Countries** | **Refs** | **Body mass (%)** | **Locomotion mode (%)** | **Coordinates (%)** |
| --- | --- | --- | --- | --- | --- | --- | --- | --- | --- |
| Home Range Area | 80416 | 1905 | 16 | 363 | 158 | 3925 | 98.39 | 100.00 | 96.84 |
| Dispersal Distance | 6868 | 1853 | 15 | 371 | 92 | 1418 | 90.99 | 99.88 | 56.10 |
| Migration Distance | 1132 | 720 | 10 | 163 | 92 | 519 | 98.32 | 100.00 | 23.14 |
| Day Range | 233 | 205 | 1 | 44 | 20 | 58 | 100.00 | 100.00 | 6.01 |
| Routine Speed | 1603 | 718 | 26 | 238 | 58 | 288 | 99.94 | 99.94 | 48.85 |
| Maximum Speed | 1955 | 729 | 21 | 216 | 32 | 242 | 99.23 | 99.95 | 2.86 |
| **Total** | **92207** | **4169** | **36** | **648** | **205** | **6234** | **97.89** | **99.98** | **89.84** |

**Figure 1.**
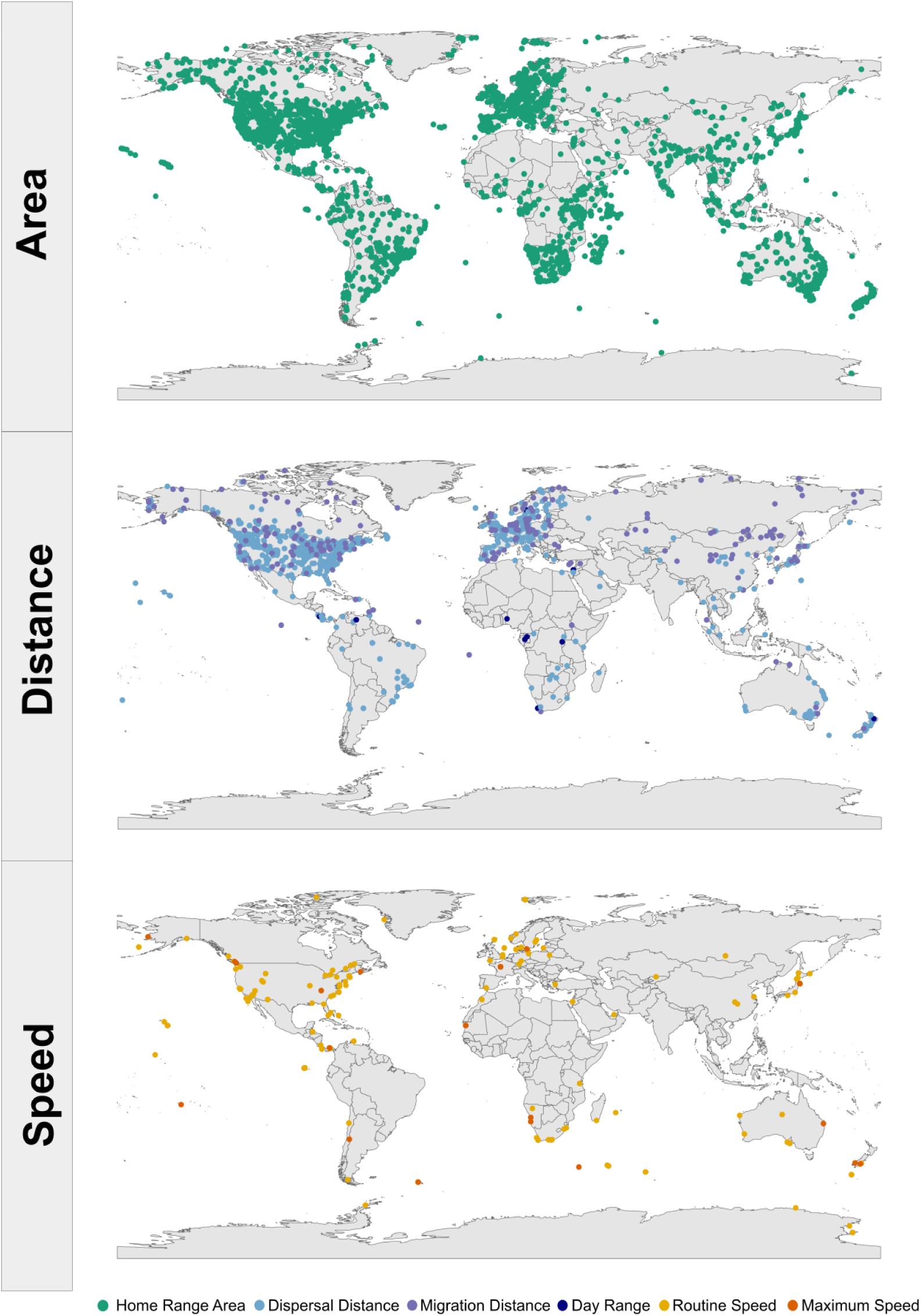
Global distribution of movement metrics in the MacroMove database. Points show the start latitude and longitude of all entries with available geographic coordinates. Movement metrics are: Home Range Area (n = 76,839; green), Dispersal Distance (n = 3,853; sky blue), Migration Distance (n = 262; purple), Day Range (n = 14; navy blue), Routine Speed (n = 374; yellow), and Maximum Speed (n = 56; orange).

**Figure 2.**
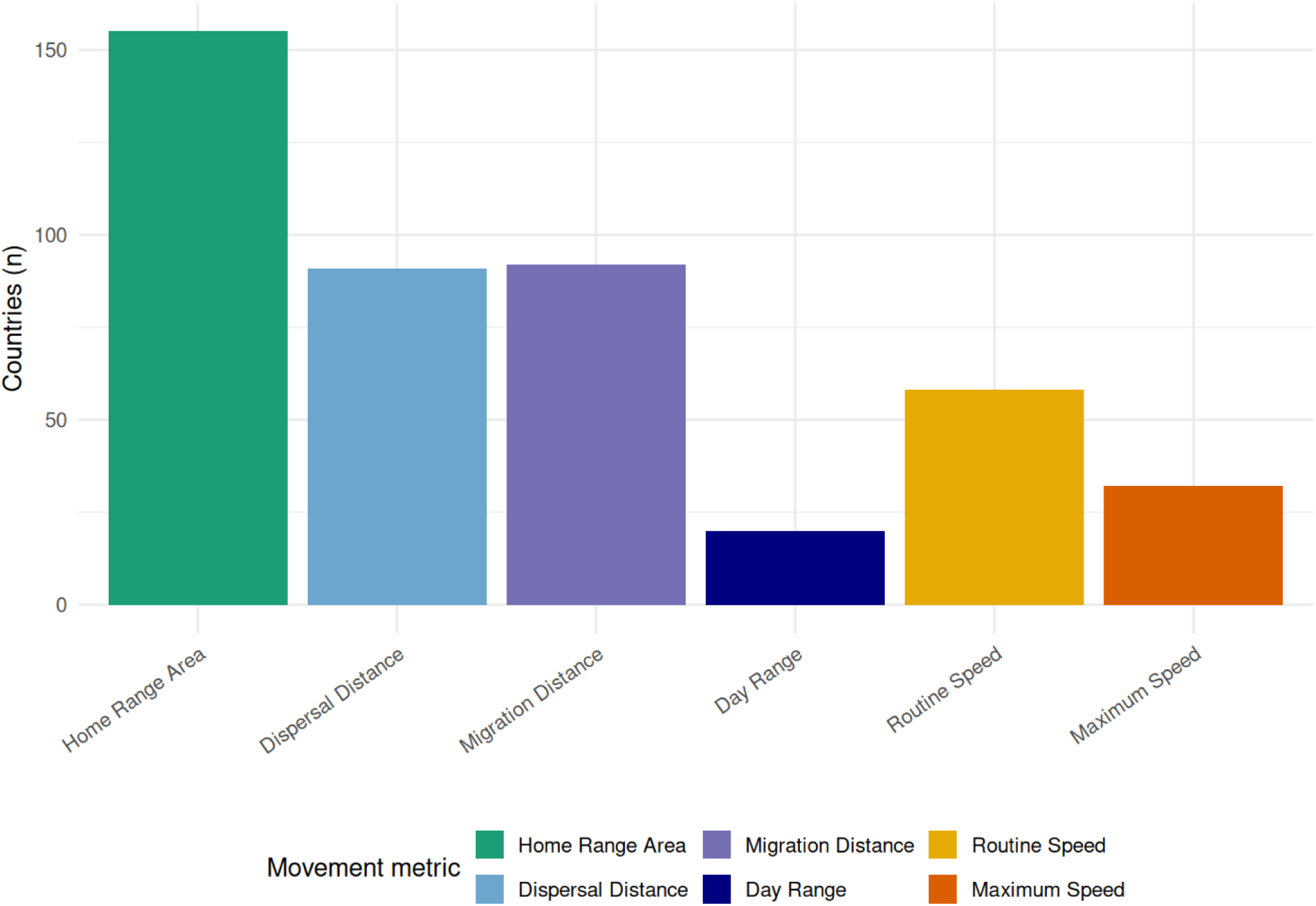
Geographic coverage of movement metrics in the MacroMove database. Bars show the number of unique countries contributing data for each movement metric. Colours indicate movement metric: Home range area (green), Dispersal distance (sky blue), Migration distance (purple), Day range (navy blue), Routine speed (yellow), and Maximum speed (orange). **Note:** Entries associated with more than one country were excluded.

**Figure 3.**
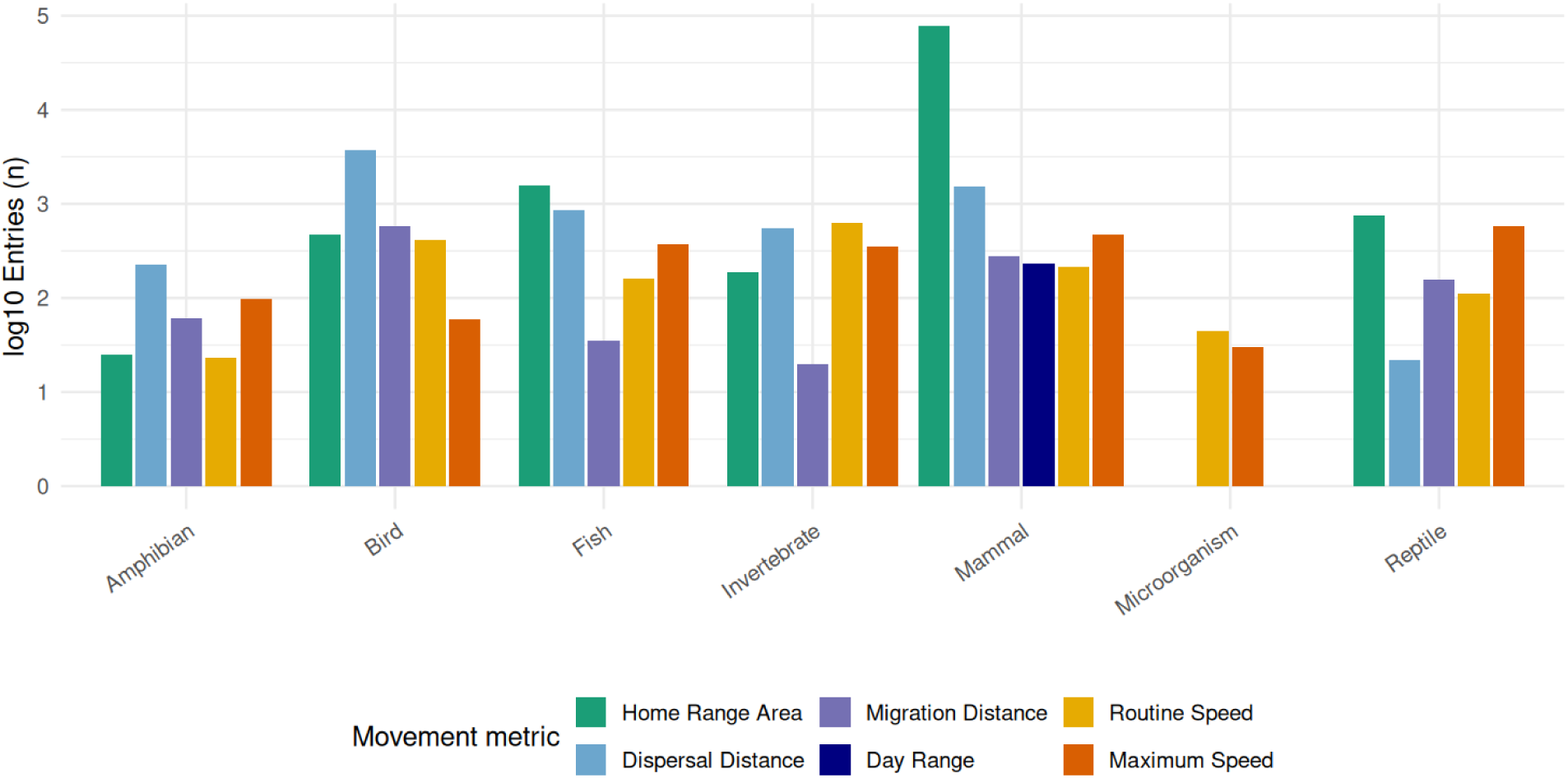
Movement observations by taxonomic category in the MacroMove database. Bars (log10 scale) show the number of entries per taxonomic category. Colours indicate movement metric: Home range area (green), Dispersal distance (sky blue), Migration distance (purple), Day range (navy blue), Routine speed (yellow), and Maximum speed (orange). **Note:** a log10 scale is applied to enable comparison across movement types with highly uneven sample sizes (see Table 1).

### Class II. Research Origin Descriptors

**A. Specific Subproject description**
  **1. Research methods** We constructed the MacroMove database by compiling information on movement processes from published, peer-reviewed meta-analyses, databases and original studies included in the meta-analyses. The meta-analyses had already aggregated movement metrics from primary sources, providing a broad foundation for cross-species comparisons across movement metrics. The literature search was focused on meta-analyses and conducted using Web of Science, applying combinations of keywords relating to movement metrics (e.g., “dispersal distance,” “migration distance,” “home range,” “movement speed,” “exploratory speed”), taxonomic scope (e.g., “organism*,” “animal*,” “vertebrate*,” “mammal*,” “bird*,” “invertebrate*,” “fish*”), and habitat type (e.g., “terrestrial,” “marine,” “aquatic,” “freshwater”), while excluding terms associated with single-species case studies. To be included, meta-analyses had to report the species’ taxonomic name, movement metric (dispersal distance, migration distance, home range area, etc.), the corresponding measurement dimension (distance, area, or speed), and a quantitative value (i.e., a numeric value rather than a range, textual description, or symbolic expression). Data were accepted from field and laboratory studies in any unit or descriptive statistic (e.g. mean, median, maximum). Additionally, we included suitable papers from any reference list of studies found through the literature search. We first identified any duplicates of original studies that were reported in multiple meta-analyses and only kept one entry for the final database. We revisited the original studies compiled by each metastudy to extract additional key variables (e.g. uncertainty value, sampling method, day/month/year, n individuals, n measurement, measurement timeframe, inference method, country, location description, coordinates, see table 3) not already included in the data files. During this step we also verified the reported values and corrected it if necessary. Additionally, we added additional movement values reported in the original studies but not in the meta-analysis (e.g. mean and median values). After this verification step, we combined the separate datasets into a single unified database. Records were imported into R for cleaning, transformation, and standardisation. Data cleaning involved standardising column names, correcting typographical errors and harmonising terminology. However, units of measurement were retained as originally reported and were not standardised across studies. To supplement missing trait information, we extracted body mass and locomotion mode from established trait databases. Data sources included COMBINE for mammals (Soria et al. 2021), AVONET for birds (Tobias et al. 2022), AnimalTraits (Herberstein et al. 2022), squamate reptiles (Slavenko et al. 2016), European amphibians (Trochet et al. 2014), marine and terrestrial vertebrates (McCauley et al. 2015), and fish and other marine fauna from FishBase and SeaLifeBase (Boettiger et al. 2023). Locomotion data reported in primary studies were retained where available. Missing values were supplemented using published databases, including Broekman et al. (2023) and Auger et al. (2024). For species not covered by these sources, locomotion mode was inferred from the corresponding family based on targeted literature and online searches. Each record includes a reference indicating the source of the locomotion information, either the primary reference, the database, or the family-based inference.
  **2. Taxonomic Data** Taxonomic names were harmonised using the R package *rgbif*, checking all species against the GBIF backbone taxonomy. Synonyms and outdated names were corrected, and accepted species names were retained. This procedure ensured consistent taxonomic resolution across MacroMove and trait databases. Across these data sources, we successfully matched between 99–100% of species names to GBIF, with nearly all sources achieving a full match. After standardisation, harmonised taxonomic and trait data were extracted from GBIF and the trait resources for subsequent data processing. Table 6 provides a detailed description of the columns in the resulting taxonomy file, which includes the original and accepted GBIF names, GBIF taxon IDs, taxonomic rank, taxonomic status, and match type.

### Class III. Data set status and accessibility

**A. Status**
  **1. Latest update:** December 2025.
  **2. Latest Archive data:** Not applicable.
  **3. Metadata status:** Metadata updated as of December 2025.
**B. Accessibility**
  **1. Storage location and medium** MacroMove is openly accessible via our Zenodo repository (XXX). All data and metadata are publicly available and may be used for research and non-commercial purposes. The repository includes version releases with specific DOIs to support reproducibility, and the dataset will be updated as needed. This project is released under the XXX license. We kindly request that users cite this work and all relevant underlying data sources upon using this dataset.
  **2. Contact people** Caitlin Wilkinson. German Centre for Integrative Biodiversity Research (iDiv) Halle-Jena-Leipzig, Theory in Biodiversity Science, Puschstraße 4, 04103, Leipzig; Friedrich-Schiller-Universitat Jena, Fürstengraben 1, 07743, Jena. Myriam R. Hirt. German Centre for Integrative Biodiversity Research (iDiv) Halle-Jena-Leipzig, Theory in Biodiversity Science,Puschstraße 4, 04103, Leipzig; Friedrich-Schiller-Universitat Jena, Fürstengraben 1, 07743, Jena. Remo Ryser. Institute for Plant Sciences (IPS), Bern, University of Bern, Bern, Switzerland.
  **3. Copyright restrictions** None. We request citation of this publication in Ecology and all relevant underlying data sources (see MacroMove_refs.csv), upon using these data.
  **4. Proprietary restrictions** None.
  **5. Costs** None.

### Class IV. Data structural descriptors

**A. Data set file**
  **1. Identity** MacroMove_db.csv MacroMove_refs.csv MacroMove_taxonomy.csv
  **2. Size** 45,842 KB 1,548 KB 5,467 KB
  **3. Format and storage mode** Comma-separated values (.csv)
  **4. Data anomalies:** If no information is available for the observation, this is indicated as ‘NA’.
**B.** Variable information

**Table 2.** Metadata for the MacroMove database (MacroMove_db.csv), describing all variables included in the main data file. Variables are listed in the order they appear in the dataset. Variables are listed in order of appearance in the dataset. Descriptions summarise each column’s content and coding conventions. Full definitions of sampling and inference methods are provided in Tables 5 and 6, respectively.

| Column Name | Storage type | Description |
| --- | --- | --- |
| Event.ID | Integer | A unique identification number for each observation. |
| Species.ID | Character | Scientific name of the species as reported in the meta-study. |
| Species.ID.gbif | Character | Scientific name of the species verified against the |
|  |  | GBIF taxonomy. |
| Taxa.category | Character | The broad taxonomic group of the species: Mammal, Bird, Invertebrate, Fish, Reptile, Amphibian, Microorganism. |
| Class.gbif | Character | Taxonomic class of the species, harmonised via GBIF. |
| Family.gbif | Character | Taxonomic family of the species, harmonised via GBIF. |
| Movement.metric | Character | The type of movement metric represented by the observation, i.e. Day range, Dispersal distance, Home range area, Maximum speed, Migration distance, or Routine speed. |
| Measurement.dimension | Character | The measurement used to quantify each movement metric, i.e. Area, Distance, or Speed. |
| Value | Real | The reported quantitative value for the movement metric. |
| Units | Character | The unit of measurement as reported in the original reference (e.g. m, km, m<sup>2</sup>, km<sup>2</sup>, km/h, m/s). |
| Statistic | Character | Descriptive statistic associated with the value (i.e. Mean, Median, Maximum, Minimum or None). None indicates when the value represents a single measurement. |
| Uncertainty.value | Real | The reported uncertainty associated with the movement measure. |
| Uncertainty.statistic | Character | The metric used to quantify the reported uncertainty value. Metrics include standard deviation (SD), standard error (SE), multiples of SE (e.g., 2×SE), confidence intervals of various levels (e.g., 80%, 90%, 95%) reported either as distance or range, interquartile range (IQR) reported as distance or range, coefficient of variation (CoV), and total range. |
| Sampling.method | Character | The empirical approach used to obtain the movement value grouped into detailed categories. Detailed definitions of each method are provided in Table 5. |
| Sampling.method.simple | Character | Simplified classification of the sampling method (e.g. Mark–recapture, Tracking, Observation, Mixed, Other, Unknown). |
| Sampling.level | Character | Indicates the hierarchical level at which the |
|  |  | movement value was measured or summarised. Values may represent a single measurement for an individual (individual-level), an average across multiple measurements for the same individual (individual-average), an average across multiple individuals of the species (species-average), an unspecified average not indicating if the value was estimated from one or multiple individuals (average), or an unknown level when the reference does not provide sufficient information. If the values were reported for a social group this is indicated by adding *group* in parentheses. |
| N.individual | Real | Number of individuals contributing to the reported value. If values are not integers the value indicates average group size. |
| N.measurement | Integer | Number of measurements contributing to the reported average value. |
| dayStart/monthStart/<br>yearStart | Integer/Character/<br>Integer | The first day/month/year from which data were collected for value calculation (starting date). May represent the first day/month/year of the study when the exact starting date for value calculation was not specified. |
| dayEnd/monthEnd/<br>yearEnd | Integer/Character/<br>Integer | The last day/month/year from which data were collected for value calculation (end date). In cases where the exact end date was not specified, the final day/month/year of the entire study was used.<br>**Note:** The difference between start and end dates does not necessarily represent the actual period over which each value was calculated. |
| Measurement.timeframe | Character | Indicates how often the movement value was observed or measured in the original study. Examples include specific time intervals (e.g. weekly, daily), fixed observation periods (e.g. 07:00–18:00), or approximate frequencies (e.g. 1–2 times per week).<br>**Note:** A category of “other” is used when the observation period does not fit these categories or was not clearly specified in the original study. When relevant, additional details such as the specific season or time of day may be provided in parentheses. |
| Inference.method | Character | Method used to calculate the home-range value. Full descriptions of each method are provided in Table 6. |
| Locomotion.mode | Character | The way in which the species moves (Climbing, Crawling, Digging, Flying, Hopping, Mixed (multiple locomotion modes), Running and Swimming).<br>**Note:** a species may exhibit different locomotion modes depending on the movement metric recorded (e.g. running for home range but flying for dispersal). |
| Body.mass.value | Real | Mass of the individual or species, as reported in the original reference, metareference or a trait database (see Body.mass.ref) |
| Body.mass.units | Character | Units of mass measurement (e.g. g, kg, log g). |
| Body.mass.sampling.level | Character | Indicates the hierarchical level at which the body mass was measured or summarised. Values may represent a single measurement for an individual (individual-level) or an average across multiple individuals of the species (species-average) |
| Sex | Character | Sex of the individuals to which the value relates (Male, Female, Both). |
| Lifestage | Character | Life stage of individuals sampled (Adult, Juvenile, Adult/Sub-adult, Juvenile/Sub-adult, Mixed, Neonate, Sub-adult). |
| Age | Character | Age of individuals, where reported. |
| Country | Character | The name of the country in which the study was conducted. If the movement process started and ended in different countries, both countries are listed. When more than two countries are involved, the value is left blank. |
| Location.description | Character | Description of the study site as reported in the original reference. |
| startLatitude/<br>startLongitude | Character | Latitude/Longitude of the study site or observation, reported in decimal degrees. When only a single location was available, it is recorded here. If multiple sites were reported, or if geographic coordinates were absent or could not be verified, this field is left blank. |
| endLatitude/<br>endLongitude | Character | Latitude/Longitude of the final location for the study or observation, when reported, converted to in decimal degrees. For migration or dispersal studies, this represents the location reached by the species. |
| Coordinates.reported | Real | Latitude and longitude of the study site or observation as reported in the original study. Coordinates were |
|  |  | verified against the study description, and corrected if they did not match the described location. If the study reported more than three locations, this field was left blank. |
| Coordinates.added | Real | Indicates the source of the decimal coordinates. 'Yes' denotes that coordinates were provided by the reference or another metareference. If coordinates were reported in an alternative coordinate system, they were converted to decimal degrees and verified using https://earth.google.com/ 'No' indicates that decimal coordinates were already provided in the metareference. |
| Original.ref | Character | The study ID assigned to the original reference. |
| Meta.ref | Character | The study ID assigned to the metareference. |
| Body.mass.ref | Character | The study ID assigned to the body mass reference. |
| Locomotion.mode.ref | Character | The study ID assigned to the locomotion mode reference. *Inferred from [Family]* indicates family-level assignment. |

**Table 3.** Metadata for the reference file used in MacroMove (MacroMove_refs.csv). Each column contains descriptive information about the source references, the verification process, and associated trait data.

| Column Name | Storage type | Description |
| --- | --- | --- |
| Study.ID | Character | The reference code assigned to the study in our dataset and reference list. |
| Citation | Character | The full citation of the reference paper. |
| DOI | Character | The DOI or online link to the reference. |

**Table 4.** Detailed descriptions of the sampling methods used in the MacroMove database (MacroMove_db.csv). Ex situ indicates deliberate manipulations by researchers, while in situ indicates natural or minimally manipulated environments.

| Sampling method simplified | Sampling method | Context | Description | Examples/ Subcategories |
| --- | --- | --- | --- | --- |
| Observation | Direct Observation | Ex situ/<br>In situ | Visual observation of animals, no instruments attached | Scan sampling, observing marked individuals |
|  | Direct Observation (Behavioural Assay) | Ex situ/<br>In situ | Visual observation of animals in response to experimental stimuli. | Escape responses, predator simulation, lab or field arenas |
|  | Instrumented Observation | Ex situ/<br>In situ | Movement recorded via instruments measuring speed, force, trajectory | Ex situ: force plates, high-speed cameras, flight mills, treadmills<br>In situ: radar guns, harmonic radar, high-speed video in field. |
|  | Indirect Observation | In situ | Movement inferred from traces | Footprints, fecal pellets, burrow spacing |
|  | Nest Box Survey | Ex situ/<br>In situ | Movement inferred from animal movement patterns using occupancy or exchange between nest boxes |  |
| Mark Recapture | Mark-Recapture | Ex situ/<br>In situ | Individuals marked and later recaptured/resighted | Banding/Ringing , PIT tags, Radioactive tags (Co60, Ta182), Elastomer, Toe/Tag, Visual marks (paint, dye) |
|  | Homing | In situ | Animals intentionally moved to measure return movement | Amphibian pond translocations, bird homing experiments |
|  | Tagging/Visual mark/Natural mark | In situ | Observing individuals marked visually | Paint, dye, metal/plastic tags, unique fur or wing patterns |
| Tracking | Radio Telemetry | Ex situ/<br>In situ | Individuals tracked remotely using VHF transmitters | Collar or tag tracking |
|  | Satellite Telemetry | Ex situ/<br>In situ | Remote tracking using GPS or ARGOS | Satellite-linked GPS tags, ARGOS PTT |
|  | Telemetry | Ex situ/<br>In situ | Other telemetry methods not included in radio or satellite | GPS-loggers, accelerometers, acoustic |
|  | Harmonic Radar | In situ | Passive tracking using radar |  |
| Other | Colonisation Experiment / Lab experiment | Ex situ | Lab/experimental manipulations | Nematode soil columns, parasitoid wasps, larval fish, experimental introductions |
|  | Powder tracking / Thread-trailing / Smoked-paper tracking | Ex situ/<br>In situ | Methods that leave visual traces | Fluorescent powder, thread spools, smoked-paper marks |
|  | Trapping / Capture | Ex situ/<br>In situ | Live traps or detection devices | Live traps, pitfall traps, camera traps |
|  | Isotope Tracing | Ex situ/<br>In situ | Tracking with radioactive or stable isotopes to infer movement | Co60, Ta182 labeling |
|  | Range Map / Survey / Census | In situ | Movement inferred from distributions or field surveys | GIS polygons, transects, point counts, habitat mapping |
|  | Experimental Rate / Introduction | In situ | Controlled releases or enclosure studies measuring movement | Salamander release studies, insect introductions, fish population spread |
|  | Beagle chases / Burrows filled with lime / Complete burrow system excavated / | In situ | Miscellaneous or specialised methods |  |
|  | Distance to nearest wetland / Movement rate |  |  |  |
| Mixed |  | Ex situ/<br>In situ | Studies that report combined methods, where multiple sampling approaches were used together. All combinations are retained as reported in the database. | Examples:<br>Radio Telemetry / Direct Observation,<br>Radio Telemetry / Trapping,<br>Trapping / Direct Observation,<br>Telemetry / Mark Recapture |
| Unknown | Unknown | NA | Method not reported or insufficient detail |  |

**Table 5.** Detailed descriptions of the inference methods used to calculate home ranges in the MacroMove database (MacroMove_db.csv). Each method represents a distinct approach to estimating animal space use.

| Inference method | Description |
| --- | --- |
| Boundary strip | External points of capture are considered centers of rectangles; the home range is the area enclosed by connecting rectangle corners. Includes inclusive and exclusive boundary strip methods. |
| Brownian Bridge | A movement-based approach where a probability density is estimated along the path between successive locations. Includes Brownian Bridge, Biased Random Bridge, and movement-based KDE methods. |
| Circle | Home range represented as a circle, e.g., based on distance from the mean location. Includes all circle methods. |
| Cluster | Home range estimated by grouping locations into clusters based on nearest-neighbor distances; clusters then merged to define area. |
| Ellipse | Home range represented by an ellipse fitted to locations; includes several ellipse-based methods. |
| Grid cell | Home range calculated by summing the area of grid cells in which the animal was located. |
| Harmonic mean | Home range contours derived from harmonic mean distances from each point to animal locations. |
| Kernel Density Estimation (KDE) | Probability density function over locations. Includes fixed, adaptive, product, time-kernel (TK), and autocorrelated KDE (AKDE) variants. |
| Local convex hulls (LoCoH) | Local convex hulls constructed for each point and its neighbors merged to estimate home range. Includes k-, r-, a-, and T-LoCoH variants. |
| Minimum Convex Polygon (MCP) | Smallest convex polygon around all locations. Includes adjusted and buffered MCP methods. |
| Minimum Area Probability | Home range estimated using probability density based on location peaks, smoothed by Fourier transformation. |
| Polygon | Polygon-based home range not specified as MCP. Includes restricted polygons and OREP methods. |
| Mixed | Home range estimated using multiple methods, e.g., averaging values from different approaches. |
| Other | Methods not fitting into the above categories (e.g., digitised polygons, multiplying river length and width). |
| Unknown | Method not reported or insufficient detail to assign category. |

**Table 6.** Taxonomic information for species included in the MacroMove database (MacroMove_taxonomy.csv). For each original species name, the table reports the source database, the GBIF-matched canonical name, GBIF usage key, taxonomic rank, taxonomic status, and match type. Hierarchical classification from phylum to species is also provided.

| Column Name | Storage type | Description |
| --- | --- | --- |
| source.ref | Character | The reference from which the species name originated. |
| original.name | Character | Original species name as provided in the database source. |
| gbif.canonical.name | Character | Canonical (accepted) name returned by GBIF. |
| gbif.usage.key | Integer | Unique GBIF identifier (usageKey) for the taxon. |
| taxonomic.rank | Character | Taxonomic rank assigned by GBIF (e.g., SPECIES, GENUS). |
| taxonomic.status | Character | Taxonomic status returned by GBIF (ACCEPTED, SYNONYM, NOT FOUND). |
| match.type | Character | Type of match performed by GBIF (e.g. EXACT, FUZZY, etc.). |
| phylum | Character | Phylum of the matched taxon according to GBIF. |
| class.name | Character | Class of the matched taxon according to GBIF. |
| family | Character | Family of the matched taxon according to GBIF. |
| genus | Character | Genus of the matched taxon according to GBIF. |
| species.name | Character | Species name assigned by GBIF (may differ from original). |

## Acknowledgements

This paper is a joint effort of the working group sMars (synthesis of movement across scales), that was kindly supported by sDiv, the Synthesis Centre of the German Centre for Integrative Biodiversity Research (iDiv) Halle-Jena-Leipzig, funded by the German Research Foundation (FZT 118). Special thanks to Stuart Butchart at BirdLife International for the unpublished bird dispersal distance data. CW and AD acknowledge the support provided by the German Research Foundation (DFG) within the research unit DynaCom (DFG, FOR 2716).

